# Isotopic reconstruction of the subsistence strategy for a Central Italian Bronze Age community (Pastena cave, 2^nd^ millennium BCE)

**DOI:** 10.1101/2022.04.21.488758

**Authors:** F. Cortese, F. De Angelis, K.F. Achino, L. Bontempo, M.R. Di Cicco, M. Gatta, C. Lubritto, L. Salari, L. Silvestri, O. Rickards, M.F. Rolfo

**Affiliations:** Department of History, Culture and Society, University of Rome Tor Vergata, Via Columbia 1, 00133, Roma, Italy; Centre of Molecular Anthropology for Ancient DNA Studies, Department of Biology, University of Rome Tor Vergata, Via della Ricerca Scientifica 1, 00133 Rome, Italy; Institute of Archaeology ZRC SAZU, Novi trg 2, 1000, Ljubljana, Slovenia; Quantitative Archaeology Lab, Department of Prehistory, Autonomous University of Barcelona, Campus UAB, 08193, Bellaterra, Spain; Fondazione Edmund Mach, Research and Innovation Centre - Traceability Unit, Via Mach 1, 38098, San Michele all’Adige, Italy; Dipartimento di Scienze e Tecnologie Ambientali, Biologiche e Farmaceutiche, Università degli Studi della Campania “Luigi Vanvitelli”; University of York, Department of Archaeology, King’s Manor, Exhibition Square, York, YO1 7EP, UK; Durham University, Department of Archaeology, Dawson Building, South Road, Durham, DH1 3LE, UK; Sovrintendenza Capitolina ai Beni Culturali, Piazza Lovatelli, 35, 00186, Rome, Italy

**Keywords:** Stable isotope analysis, Bronze Age, Central Italy, Crop management, Animal husbandry, Diet reconstruction

## Abstract

The Pastena cave is located in central Italy and its best-preserved sector is Grotticella W2, which is dated radiometrically to the Early-Middle Bronze Age. The aim of this paper is to explore human diet, animal husbandry, and plant management analyzing the findings there discovered. Carbon and nitrogen stable isotope analysis have been carried out on 40 charred seeds, 6 faunal remains and 4 human specimens, investigating the whole bio-archaeological samples available. To the best of our knowledge, this is one of the first papers presenting stable isotope analysis on carpological remains dated to the Italian Early-Middle Bronze Age. The obtained results are consistent with a diet based on terrestrial protein, mainly on plants and secondly on meat and animal products. The data suggest that plants, especially broad beans, were partially subjected to human management, while livestock was managed through different husbandry strategies. The cooperation between archaeological studies and molecular analysis allows us to contribute to clarify the economic strategies for a Central Italian community in a scenario that is still poor in published data.

## Introduction

The Bronze Age (BA) is one of the crucial periods of late Italian Prehistory (Bietti Sestieri 2015). Even though this phase presents different characteristics and chronologies in Europe (Fokkens, Harding 2013), in the Italian peninsula, it is assumed that its chronological boundaries dated around 2300-950 BCE (Peroni 1996; Bietti Sestieri 2015). Despite the plethora of archeological surveys (e.g., Cocchi Genick, 1995; Barfield 2007; Guidi, Rosini 2019; Skeates et al. 2021), the bioarcheological ones are still sparse and refer to sites scattered throughout the Peninsula, with a lack of data relating to Central-South Italy. So far, the main object of our study is the Pastena cave, which returned traces of human frequentation starting from the Late Neolithic to the Middle Bronze Age (MBA) (Guareschi & Morandini, 1943; Biddittu 1987; Biddittu et al. 2007; Angle et al. 2010; Rolfo et al. 2021). Specifically, we focus on the BA bioarcheological findings, which could act as significant pieces of evidence for supporting hypotheses related to complex dynamics occurring in the Central Italian BA, contributing to the claimed need for denser and timely-resolved evidence to assess the spread of BA cultures in Italy (Saupe et al. 2021; Romboni et al. 2022).

Indeed, the BA is a period of massive changes in technological, cultural, social and economic settings for human communities (Fokkens, Harding 2013). The spread of bronze metallurgy is the most important technological innovation, and its processing required specialized roles leading to the rise of social complexity. Indeed, as the bronze items were often considered prestige goods and personal ornaments, their availability reflected the social stratification of the communities, which was supposed to be reflected, in turn, by the discovery of these items as goods in the burial grounds (Bettelli 2006; Fokkens, Harding 2013).

A complex territorial organization also characterizes BA through the development of hierarchical residential systems, which again suggest a complex social organization: nucleated settlements, fortified villages, pile-dwellings, and terramare hosted a growing number of inhabitants (Bettelli 2006; Fokkens, Harding 2013). Despite the increasing establishment of settlements, the discovery of exotic items and recent stable isotope analysis on human bones suggest a rise in both individual and artifact mobility in the BA (e.g., Knipper et al. 2017; Cavazzuti et al. 2019; Cavazzuti et al. 2021). Goods and people were involved in complex exchange networks (Blake 2014; Cavazzuti et al. 2019), whose effects could be identified in heterogeneous lifestyles and bio-cultural characteristics (Fokkens, Harding 2013; Bietti Sestieri 2015).

A multifaceted funerary scenario also reflected the heterogenous dwelling strategy. Multiple regional specificities could be outlined across the Italian peninsula, mainly concerning pit graves, mounds, or artificial and natural cave burials, where the last were often used for ritual purposes (Bettelli 2006; Whitehouse 2007; Minniti 2012; Bietti Sestieri 2015).

Indeed, most of the knowledge for BA in Central Italy came from karst environments, as numerous caves hosting BA evidence characterize this area (Sestini 1934; Rittatore 1951a, 1951b; Cocchi Genick 1995; Cavanna 2007; Alessandri et al. 2021; Rolfo et al. 2021).

So far, the analysis of biological remains from multiple sites complemented the archeological findings and supported a comprehensive understanding of the complex dynamics occurring in the Central Italian BA.

The carbon and nitrogen stable isotope analysis on bio-archaeological remains has been steadily considered able to identify the dietary habits, and in turn, the economy and social behavior of past communities (e.g., Shoeninger, DeNiro 1984; Katzenberg 2008; Fontanals-coll et al. 2015; Goude et al. 2016; De Angelis et al. 2019; De Angelis et al. 2020; Varalli et al. 2021; Romboni et al. 2022). Briefly, the collagen from bones could be extracted and its carbon and nitrogen composition would relate to the individual dietary landscape. Specifically, the carbon stable isotopes (^12^C and ^13^C) ratios in bone collagen account for terrestrial or marine protein sources, though they also mark for distinguishing the photosynthetic pathways of the exploited plants. Conversely, the analyses of nitrogen stable isotopes (^14^N and ^15^N) ratios allow for disentangling the trophic level, generally assigning species to herbivores, omnivores and carnivores (De Niro 1985; Ambrose 1990; Hedges et al. 2007).

Human dietary habits are one of the most retained markers of the people and societies’ cultural identity (De Angelis et al. 2019) and have been extensively dissected through stable isotope analysis across the Italian BA communities (e.g., Varalli et al. 2015; Tafuri et al. 2018; Arena et al. 2020; Romboni et al. 2022). Indeed, these findings are consistent with the archeological and archeozoological data, suggesting that the BA human groups had a mixed economy, mainly based on agriculture and breeding practices tightly related to the local ecosystem. Conversely, wild gathering through hunting and fishing was less practiced (De Grossi Mazzorin 1995; Cazzella 2009; Cattani, Marchesini 2010; Maini 2010; Rolfo et al. 2013; Salari, Tagliacozzo 2019).

However, new crops spread throughout the European continental areas during the BA (Miller et al. 2016). C_4_ plants (e.g., the common millet, *Panicum miliaceum*, or broomcorn millet) appeared consistently in Europe in the late 3^rd^ millennium BC, to become established during the second half of the 2^nd^ millennium BC (Stevens et al. 2016; Valamoti 2016). These plants are well suited to an arid ecology. As fast-growing, warm-season crops, their cultivation was probably started to be cultivated to reduce agricultural risk as a low-investment rain-fed crop (Stevens et al. 2016). The impact of these plants on the Italian BA diet has been previously outlined even through carbon and nitrogen stable isotope analysis (Tafuri et al. 2009, 2018; Varalli et al. 2015; Romboni et al. 2022), supporting the role of the human population dispersal across the peninsula (Saupe et al. 2021; Romboni et al. 2022) for the dissemination of these resources. Evidence of C_4_ plants consumption was also reported for previous periods in Italy (Mariotti-Lippi et al. 2017; Nava et al. 2021). However, whether these C_4_ plants were gathered as wild plants or served as animal fodder rather than extensively cultivated and consumed by people is not fully understood. Despite the local or occasional consumption of these crops, the diet of people living in the Italian peninsula during the BA seems to have primarily consisted of C_3_ plants to be complemented by a moderate amount of animal protein (Arena et al. 2020). Indeed, the prevailing horticultural regime for Peninsular Italian communities had long depended on wheat (*Triticum sp*.) and barley (*Hordeum vulgare*), whose charred remains were often recovered in archeological surveys (e.g., Tongiorgi 1950; Guidi 1991-92; Pacciarelli 1997; Guidi, Rosini 2019). These grains were widespread in the BA contexts testifying that these plant species were broadly available. Indeed, they could be cultivated, even accounting for farming strategies implemented in productive economies to sustain their harvesting in geographical areas characterized by moist winters and meager summer precipitation.

Remarkably, other than outlining the direct trophic relationship, the stable isotope analysis could also be meaningful for disentangling these horticulture practices (Ferrio et al. 2005; Bogaard et al. 2007, 2013; Flohr et al. 2011; Fraser et al. 2011; Kanstrup et al. 2014; Szpack et al. 2014; Treasure et al. 2016; Gröcke et al. 2021). The analysis of the plant remains has often supported the archeological identification of horticultural practices, which were critical for developing farming strategies. Specifically, fertilization and irrigation practices, recognizable with changes in carbon and nitrogen stable isotopes ratios, are two of the most significant technological improvements for boosting soil exploitation, supporting both sedentary lifestyles and the market economy (Wallace et al. 2015; Styring et al. 2016; Gron et al. 2017, 2020; Knipper et al. 2020; Mnich et al. 2020; Varalli et al. 2021).

In that perspective, the stable isotope analysis of the faunal remains, which is generally carried out to establish the local isotopic variability, can also reveal, in turn, breeding strategies and the use of specific agricultural practices (Goude et al. 2016). Accordingly, this work aims at contributing to the reconstruction of the economic identity of people buried in the Pastena cave (Frosinone, Latium) during the first half of the BA. The results were put in the context of recent genomic and isotopic studies, which provided suggestions for the population dynamics (Saupe et al. 2021; Romboni et al. 2022) and technological improvements (Varalli et al. 2015) running in Central Italian Early and Middle BA (Fig. 1A).

**Fig. 1:**
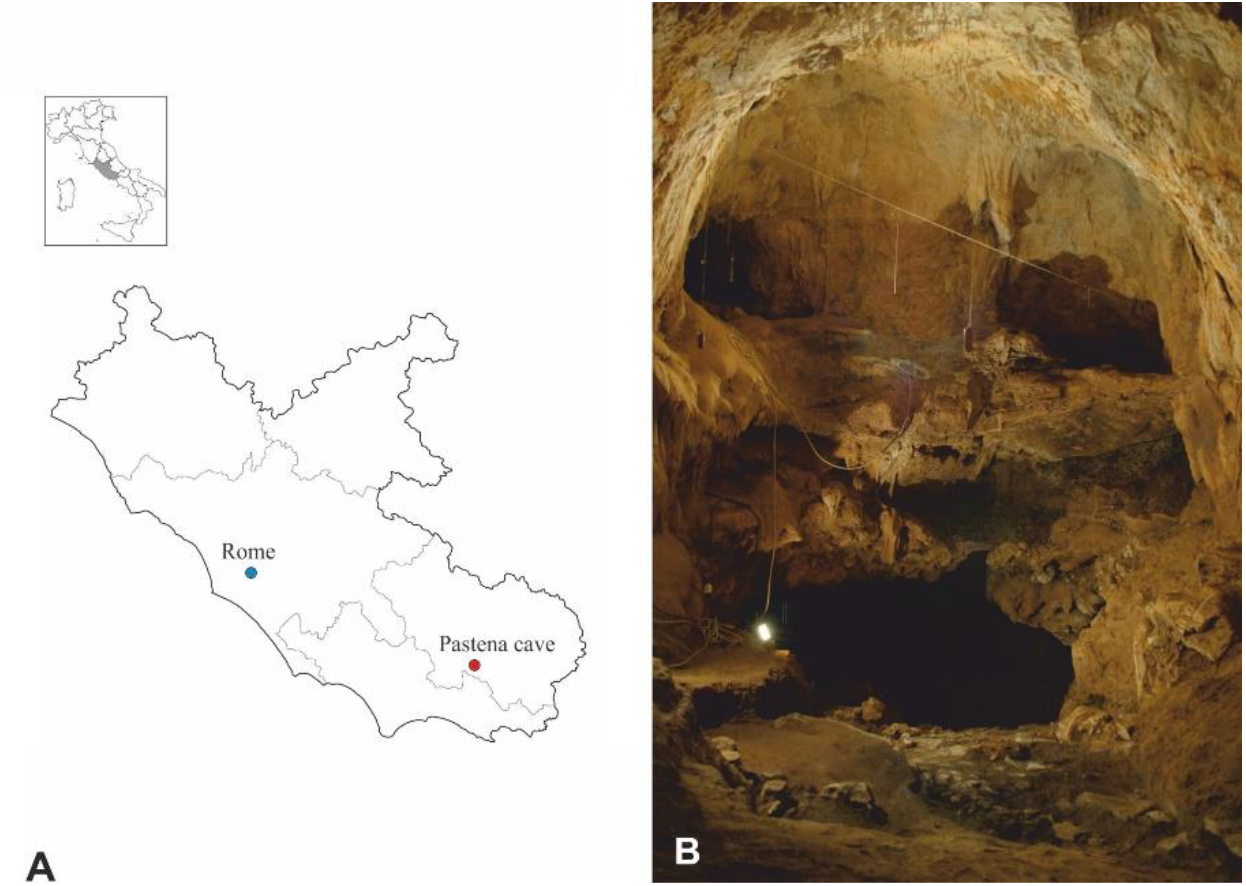
A) Pastena Cave in the Italian peninsula; B) East view of Grotticella W2 with its upper terrace.

### Archaeological context

The Pastena cave (41°29′48.77″N, 13°29′21.91″E) is situated approximately 196 meters a.s.l. in a karst valley. The entrance presents a majestic chamber, which features a seasonal stream of the Rio Mastro Creek. The first excavations were performed in the first half of the 20^th^ century (Guareschi, Morandini 1943), and subsequently, surveys and excavations were carried out in several areas of the cave (Biddittu 1987; Biddittu et al. 2007; Angle et al. 2010), discovering archeological findings ranging between the Late Neolithic and the Middle Bronze Age.

The investigation focusing on Grotticella W2 (GW2, Fig.1B) started in 2006 and subsequently was carried out thoroughly between 2012 and 2018 (Angle et al. 2014; Silvestri et al. 2019; Rolfo et al. 2021). This area is a small room opening in the west sector of the cave’s entrance hall and it is the only area not affected by post-depositional events. The archaeological deposit has been thoroughly investigated, and the radiocarbon dating on seeds and human remains already suggested that the deposit was mono-phasic and related to the end of the Early Bronze Age (EBA) up to the Middle Bronze Age (MBA) (Rolfo et al. 2021).

Six main layers including multiple stratigraphic units and archaeological structures, such as hearths, reddened areas, stone paving and pits, as well as several archaeological finds, including pottery and personal ornaments have been identified in GW2, suggesting its intensive use related to ritual and funerary practices (Silvestri et al. 2019; Rolfo et al. 2021).

The artifacts were found intermingled along with seeds and bones. Thousands of burnt carpological remains were recovered: botanical visual analyses revealed that they mostly pertained to *Vicia faba*, while cereals (*Triticum monococcum/dicoccum, Hordeum vulgare, Triticum aestivum*) account for the residual fraction.

A few faunal remains were found in GW2 (about 100 finds): they are primarily domesticated mammals, especially sheep/goats and pigs, while wild species were rare (Silvestri et al. 2019).

Humans are represented by a minimum number of individuals (MNI) of 4, belonging to different age classes, including osteologically immature ones. The human bones mostly consist of distal limb segments, except for a mandible, some teeth, and a few long bones, which were helpful in estimating age at death and MNI (Rolfo et al. 2021).

## Materials and Methods

We performed stable isotope analyses on four human bones, eight faunal remains belonging to domesticated animals (*Ovis aries* vel *Capra hircus, Sus domesticus*, and *Canis familiaris*), and forty burnt seeds (*Vicia faba, Triticum monococcum/dicoccum, Triticum aestivum*, and *Hordeum vulgare*) (Silvestri et al. 2019; Rolfo et al. 2021).

Carbon and nitrogen stable isotope analysis is commonly used to reconstruct past dietary habits (DeNiro 1985; Ambrose 1990; Unkovich et al. 2001; Marshall et al. 2007). The method is based on estimating the ratios of ^13^C/^12^C and ^15^N/^14^N, detectable from the collagen, which reflect the diet of the last decades of life (Ambrose 1990; Hedges et al. 2007). The relative abundance of ^13^C and ^15^N is expressed in per mill (‰) through δ notation (respectively δ^13^C and δ^15^N) with respect to international standards - Vienna Pee Dee Belemnite (VDB) for δ^13^C and atmospheric nitrogen (AIR) for δ^15^N - according to the relationship: δiE = (iRSA-iRREF) / iRREF, where *i* is the mass number of the heavier isotope of the E element; RSA is the isotope ratio of the sample; and RREF is the relevant internationally recognized reference material (Mariotti 1983).

The set of carbon and nitrogen values reflects the trophic position of an individual between the available resources. Briefly, the trophic relations are based on a direct correlation between prey and predator: the latter is higher than its game’s value (DeNiro 1985). Previous studies accounted for an average offset of about 1‰ in δ^13^C and 3-5‰ in δ^15^N values observed between two contiguous trophic levels (DeNiro 1985).

The δ^13^C ratio is suitable to determine the consumption of C_3_ (typical of temperate environments) and C_4_ plants (mainly from arid ecosystems), as well as marine and freshwater intake (Ambrose, Norr 1993; Krigbaum 2003; Marshall et al. 2007; Katzenberg 2008). Conversely, the δ^15^N provides information related to the trophic levels, basically relying on protein intake (Shoeninger, DeNiro 1984; Keegan 1989; Unkovich et al. 2001).

Collagen was extracted from cortical bones for humans and faunal remains. Samples were prepared at the Centre of Molecular Anthropology for Ancient DNA Studies of the University of Rome “Tor Vergata”, following Longin’s protocol (1971), modified by Brown et al. (1988). The cortical bone was collected and cleaned by surface abrasion. The extraction protocol was performed on about 800mg of bone and a modern bovine sample was used as a reference. Samples were demineralized in HCl 0.6M, rinsed with bi-distilled water, and finally gelatinized in HCl 0.001 M. An ultrafiltration step through 30kDa Amicon Ultra-4 Centrifugal Filter Unites with Ultracel membranes (Millipore) was executed to maximize the collagen concentration and the obtained collagen was lyophilized. Carbon and nitrogen stable isotope ratios were measured in a single run on a Delta V Advantage isotope ratio mass spectrometer coupled to a Flash 1112 Elemental Analyser via a Conflow III interface (Thermo Scientific Milan, Italy) at Dipartimento di Scienze e Tecnologie Ambientali, Biologiche e Farmaceutiche – Università degli Studi della Campania Luigi Vanvitelli and on a Delta Plus XP isotope ratio mass spectrometer coupled with a Flash 1112 Elemental Analyser via a Conflow IV interface (Thermo Scientific Milan, Italy) at Food Quality and Nutrition Department, Traceability Unit – Fondazione Edmund Mach. The batches compared and no deviances were detected. The samples were compared against established criteria to ascertain the percentages of carbon and nitrogen, atomic C/N ratios and collagen yields (DeNiro 1985; Ambrose 1990; van Klinken 1999); analytical precision is ± 0.2‰ for δ^13^C, reported with respect to the PDB standard, and ± 0.3‰ for δ^15^N, reported with respect to AIR. To assess the preservation state of the extracted collagen, we considered carbon and nitrogen contents between 15-51% and 5-18%, respectively, and C/N ratios within the range of 2.9 to 3.6 (DeNiro 1985; Ambrose 1990).

An established protocol for stable isotope analysis is not available for seeds. Different methods have been applied: acid-base-acid pre-treatment protocol (Bogaard et al. 2013; Fraser et al. 2013b) and crushing and powdering raw seeds (Heaton et al. 2009; Gron et al. 2017, 2020) were the most applied approaches in recent attempts. Considering the results obtained by Vaiglova et al. (2014), we crashed the seeds in a mortar and the powder was inlet into the isotope ratio mass spectrometer. No consensual criteria exist to assess the preservation of charred seeds and chaff; thus, we retained all the samples returning positive yields (Vaiglova et al. 2014).

Faunal and human data have been evaluated through the linear mixing model proposed by Fraser et al. (2013a). As described by the authors, this model uses the midpoint and the offsets between consecutive trophic levels - an average of 4‰ for δ^15^N - to identify the effect of predators on their prey. Accordingly, we implemented different models based on plants and faunal values determined in the present paper to calculate the theoretical endpoints for terrestrial resources consumption to evaluate the intake of domesticated crops in faunal remains and the animal protein consumption in the human diet.

Furthermore, we elaborated on the Bayesian mixed models developed through FRUITS v3.1 (Fernandes et al. 2014). The software provides probabilistic quantification of dietary inputs and incorporates food macronutrients, elemental composition and isotopic composition in its calculation. The average isotopic values of humans were used as consumer data and all calculated values were input into FRUITS with instrumental uncertainties of 0.1‰ for carbon and 0.2‰ for nitrogen. Four resources (cereals, broad beans, fauna, freshwater fish) have been identified as putative food groups in the model. We used the data from local resources for the first three categories, while freshwater fish values are based on published values (Varalli et al. 2021). The carbon and nitrogen offset/weight ratios are based on Hedges, Reynard (2007), Fernandes et al. (2015) and Styring et al. (2017); and the estimation of source values and the amount of energy and proteins in each food groups were calculated following Fernandes et al. (2015).

Seeds watering status has been established through carbon values. In order to consider CO_2_ fluctuations in Holocene, Δ^13^C values have been used to calculate water availability, comparing archaeological plants data over time (http://web.udl.es/usuaris/x3845331/AIRCO2_LOESS.xls, Ferrio et al. 2005). Δ^13^C values reflect plants’ water status during their life cycle, which was influenced both by natural precipitation and human management (Wallace et al. 2015). Wallace et al. (2015) provide the largest collections of Δ^13^C values for the archaeological crop, to be used as patterns for the watering status of ancient plants. Following Ferrio et al. (2005), we calculated the plant Δ^13^C from δ^13^C_air_ and plant carbon isotope composition as described by Farquhar et al. (1982) (See Suppl. Mat.).

The estimation of manuring rates was addressed through the comparison with methodological approaches (e.g., Bogaard et al. 2007; Styring et al. 2016; Treasure et al. 2016) and values obtained from archaeological contexts (e.g., Kanstrup et al. 2014; Bogaard et al. 2013), showing that cereals not subjected to fertilization have a baseline δ^15^N around 2.5‰. Conversely, manured grains (about 35 t/ha compost applied annually) present nitrogen values equal to or greater than 6‰. Intermediate δ^15^N values between 2.5‰ and 6‰ can be related to different scenarios (Treasure et al. 2016). Differently, legumes naturally present δ^15^N values of 0‰, and only an intensive fertilization with animal cages (>70 t/ha) can increase nitrogen values to reach +3‰ (Treasure et al. 2016).

## Results

Carbon and nitrogen isotope values of seeds, animals, and humans are listed in Table 1 and shown in Fig. 2. Isotope data were successfully obtained for all the materials, but two samples of *Sus domesticus* were excluded due to their C/N ratios outside the range 2.9-3.6 (DeNiro 1985).

**Tab. 1.**
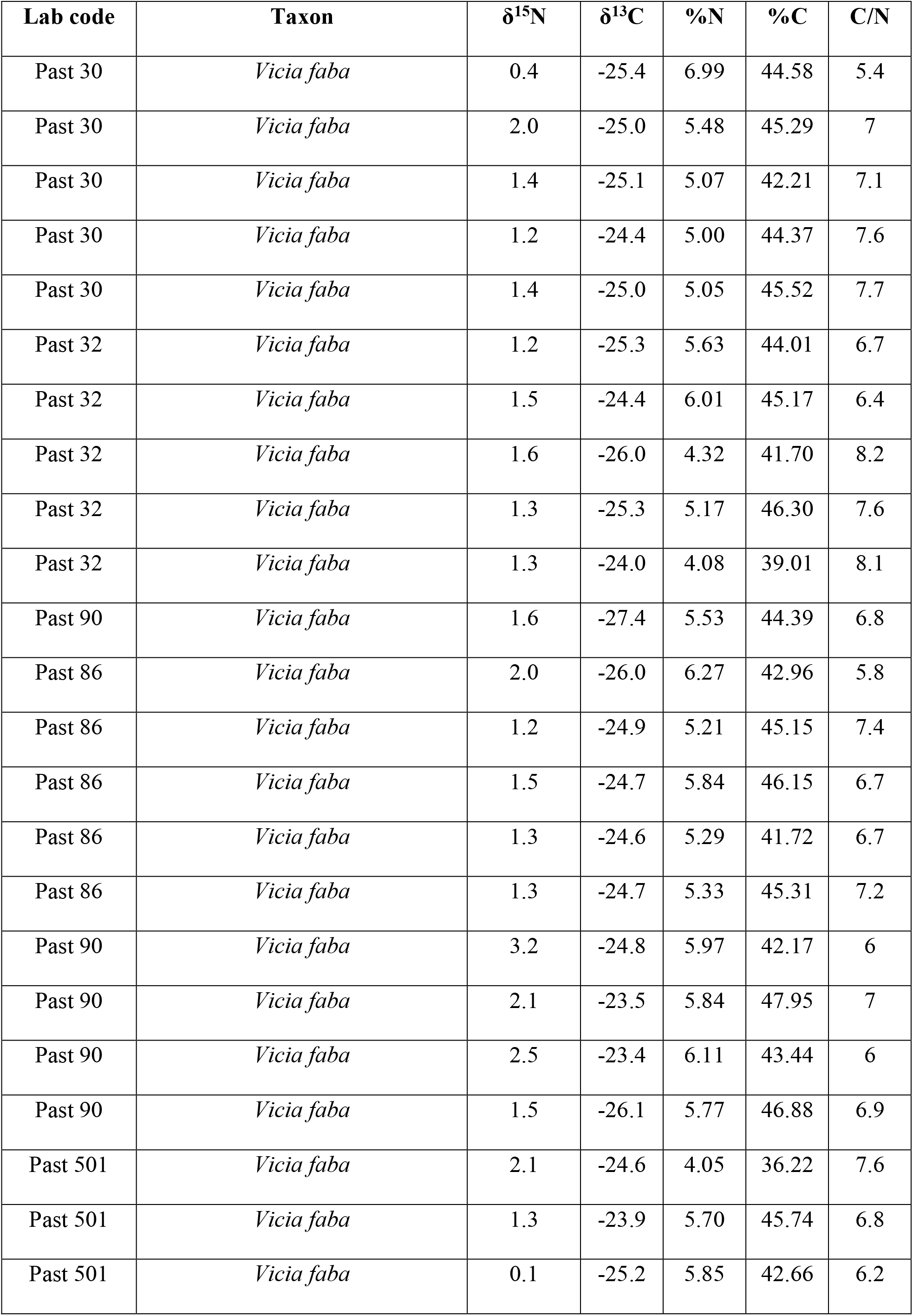
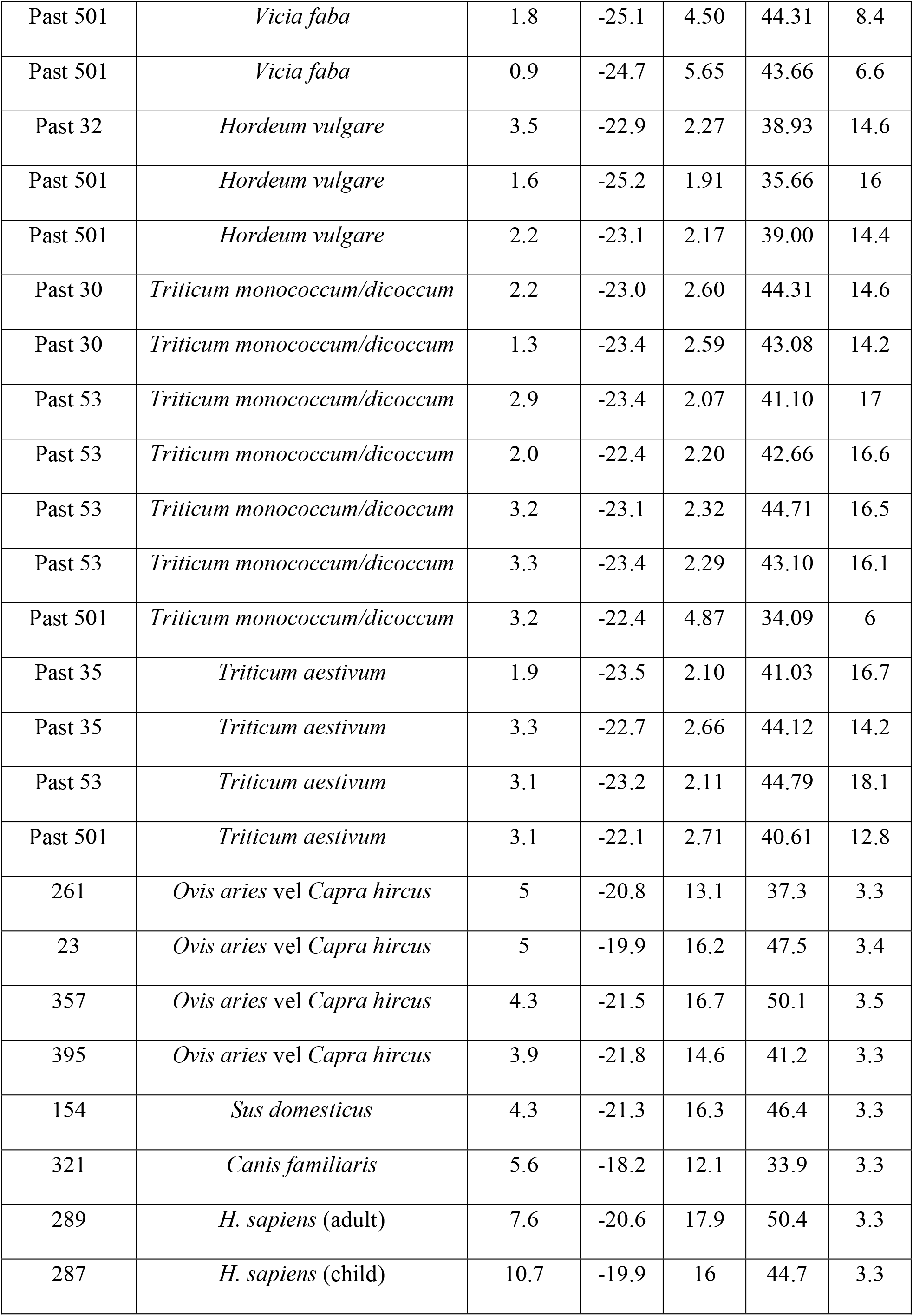
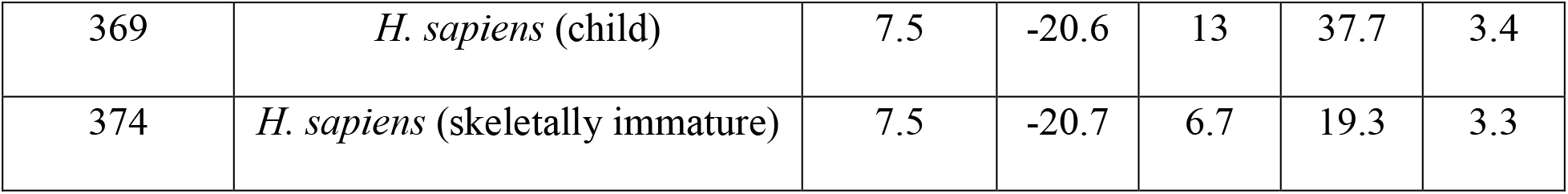
Stable isotope analysis results, and collagen quality control indicators for plants, fauna and humans. See Suppl. Mat. for the estimation of δ^15^N considering the charring effect.

| Lab code | Taxon | $\delta^{15}\text{N}$ | $\delta^{13}\text{C}$ | %N | %C | C/N |
| --- | --- | --- | --- | --- | --- | --- |
| Past 30 | *Vicia faba* | 0.4 | -25.4 | 6.99 | 44.58 | 5.4 |
| Past 30 | *Vicia faba* | 2.0 | -25.0 | 5.48 | 45.29 | 7 |
| Past 30 | *Vicia faba* | 1.4 | -25.1 | 5.07 | 42.21 | 7.1 |
| Past 30 | *Vicia faba* | 1.2 | -24.4 | 5.00 | 44.37 | 7.6 |
| Past 30 | *Vicia faba* | 1.4 | -25.0 | 5.05 | 45.52 | 7.7 |
| Past 32 | *Vicia faba* | 1.2 | -25.3 | 5.63 | 44.01 | 6.7 |
| Past 32 | *Vicia faba* | 1.5 | -24.4 | 6.01 | 45.17 | 6.4 |
| Past 32 | *Vicia faba* | 1.6 | -26.0 | 4.32 | 41.70 | 8.2 |
| Past 32 | *Vicia faba* | 1.3 | -25.3 | 5.17 | 46.30 | 7.6 |
| Past 32 | *Vicia faba* | 1.3 | -24.0 | 4.08 | 39.01 | 8.1 |
| Past 90 | *Vicia faba* | 1.6 | -27.4 | 5.53 | 44.39 | 6.8 |
| Past 86 | *Vicia faba* | 2.0 | -26.0 | 6.27 | 42.96 | 5.8 |
| Past 86 | *Vicia faba* | 1.2 | -24.9 | 5.21 | 45.15 | 7.4 |
| Past 86 | *Vicia faba* | 1.5 | -24.7 | 5.84 | 46.15 | 6.7 |
| Past 86 | *Vicia faba* | 1.3 | -24.6 | 5.29 | 41.72 | 6.7 |
| Past 86 | *Vicia faba* | 1.3 | -24.7 | 5.33 | 45.31 | 7.2 |
| Past 90 | *Vicia faba* | 3.2 | -24.8 | 5.97 | 42.17 | 6 |
| Past 90 | *Vicia faba* | 2.1 | -23.5 | 5.84 | 47.95 | 7 |
| Past 90 | *Vicia faba* | 2.5 | -23.4 | 6.11 | 43.44 | 6 |
| Past 90 | *Vicia faba* | 1.5 | -26.1 | 5.77 | 46.88 | 6.9 |
| Past 501 | *Vicia faba* | 2.1 | -24.6 | 4.05 | 36.22 | 7.6 |
| Past 501 | *Vicia faba* | 1.3 | -23.9 | 5.70 | 45.74 | 6.8 |
| Past 501 | *Vicia faba* | 0.1 | -25.2 | 5.85 | 42.66 | 6.2 |
| Past 501 | *Vicia faba* | 1.8 | -25.1 | 4.50 | 44.31 | 8.4 |
| Past 501 | *Vicia faba* | 0.9 | -24.7 | 5.65 | 43.66 | 6.6 |
| Past 32 | *Hordeum vulgare* | 3.5 | -22.9 | 2.27 | 38.93 | 14.6 |
| Past 501 | *Hordeum vulgare* | 1.6 | -25.2 | 1.91 | 35.66 | 16 |
| Past 501 | *Hordeum vulgare* | 2.2 | -23.1 | 2.17 | 39.00 | 14.4 |
| Past 30 | *Triticum monococcum/dicoccum* | 2.2 | -23.0 | 2.60 | 44.31 | 14.6 |
| Past 30 | *Triticum monococcum/dicoccum* | 1.3 | -23.4 | 2.59 | 43.08 | 14.2 |
| Past 53 | *Triticum monococcum/dicoccum* | 2.9 | -23.4 | 2.07 | 41.10 | 17 |
| Past 53 | *Triticum monococcum/dicoccum* | 2.0 | -22.4 | 2.20 | 42.66 | 16.6 |
| Past 53 | *Triticum monococcum/dicoccum* | 3.2 | -23.1 | 2.32 | 44.71 | 16.5 |
| Past 53 | *Triticum monococcum/dicoccum* | 3.3 | -23.4 | 2.29 | 43.10 | 16.1 |
| Past 501 | *Triticum monococcum/dicoccum* | 3.2 | -22.4 | 4.87 | 34.09 | 6 |
| Past 35 | *Triticum aestivum* | 1.9 | -23.5 | 2.10 | 41.03 | 16.7 |
| Past 35 | *Triticum aestivum* | 3.3 | -22.7 | 2.66 | 44.12 | 14.2 |
| Past 53 | *Triticum aestivum* | 3.1 | -23.2 | 2.11 | 44.79 | 18.1 |
| Past 501 | *Triticum aestivum* | 3.1 | -22.1 | 2.71 | 40.61 | 12.8 |
| 261 | *Ovis aries* vel *Capra hircus* | 5 | -20.8 | 13.1 | 37.3 | 3.3 |
| 23 | *Ovis aries* vel *Capra hircus* | 5 | -19.9 | 16.2 | 47.5 | 3.4 |
| 357 | *Ovis aries* vel *Capra hircus* | 4.3 | -21.5 | 16.7 | 50.1 | 3.5 |
| 395 | *Ovis aries* vel *Capra hircus* | 3.9 | -21.8 | 14.6 | 41.2 | 3.3 |
| 154 | *Sus domesticus* | 4.3 | -21.3 | 16.3 | 46.4 | 3.3 |
| 321 | *Canis familiaris* | 5.6 | -18.2 | 12.1 | 33.9 | 3.3 |
| 289 | *H. sapiens* (adult) | 7.6 | -20.6 | 17.9 | 50.4 | 3.3 |
| 287 | *H. sapiens* (child) | 10.7 | -19.9 | 16 | 44.7 | 3.3 |
| 369 | *H. sapiens* (child) | 7.5 | -20.6 | 13 | 37.7 | 3.4 |
| 374 | *H. sapiens* (skeletally immature) | 7.5 | -20.7 | 6.7 | 19.3 | 3.3 |

**Fig. 2:**
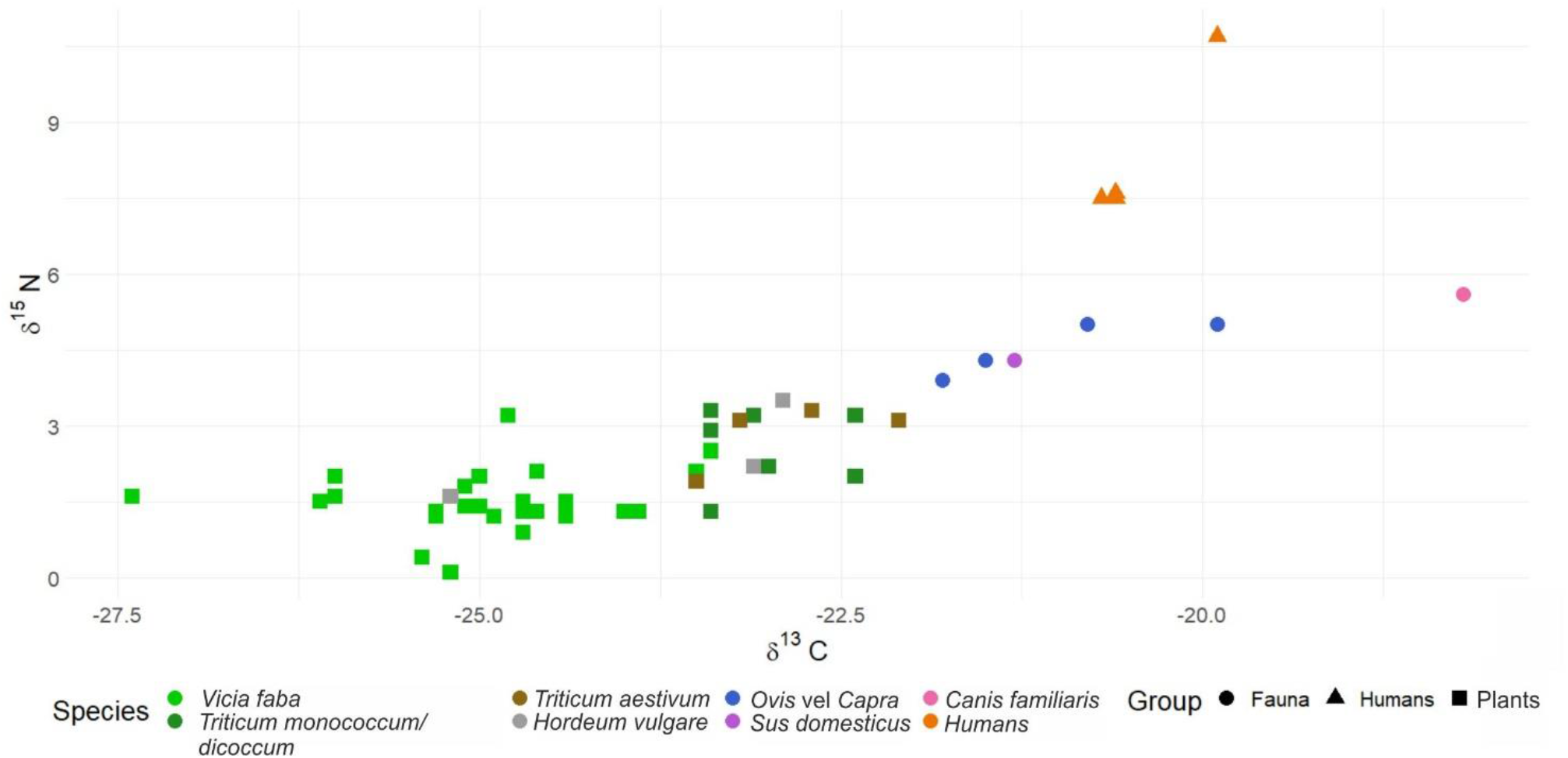
Isotope data dispersion for seeds, faunal remains and humans from the Pastena cave.

A whole sample of 40 seeds, constituted by broad beans, emmer, barley, and wheat, was also characterized. The isotopic ratios of the C_3_ plants range from 0.1‰ to 5.2‰ for δ^15^N (median= 1.7‰; IQR= 1.60), and −22.1‰ to −27.4‰ for δ^13^C (median= −24.5‰; IQR= 1.80).

For Δ^13^C, calculated according to Ferrio et al. (2005), values varied between 17.5‰ and 21.6‰ for legumes and between 16.1‰ and 19.3‰ for cereals, indicating that these crops are grown under good hydric conditions. The considerable variation in legumes and grains suggests that they were cultivated on soils receiving variable water inputs and/or varying water retention capacity. Seeds values have been tested through the Kolmogorov-Smirnov test: cereals do not present significant differences (*p*>0.05), possibly due to the small sample size. Legumes and cereals appear significantly different in both carbon (*p*=5.65E-07) and nitrogen values (*p*=2.80E-04). Considering the lack of outlier in these two groups and consistent with archaeological information, we consider the values representative of local produce. The isotopic markers make reliable that agricultural practices, such as manuring or watering, were present, even though they do not appear to have been as extensive as we expected, considering the vast quantities of seeds in the deposit.

The domesticated animals show δ^13^C values ranging between −20.8‰ and −18.2‰ (IQR= 1.60), and 3.9‰ and 5.6‰ (IQR= 0.65) for δ^15^N. Their values suggest that herbivores were left to graze.

We tried to confirm the hypothesis mentioned above by evaluating the fraction of plants from the total intake and realizing two models based on the δ^15^N. The first one (Fig. 3A) is based on the mean value of *Vicia faba*, which is the starting point to estimate the intake of these crops in herbivores’ diet, as they should rely upon a +4‰ step. The returned consumption of legumes proposed by that model is very high (about 80%). The second model (Fig. 3B) takes into account the mean value of cereals: the result suggests a low intake of these plants (about 40%).

**Fig. 3:**
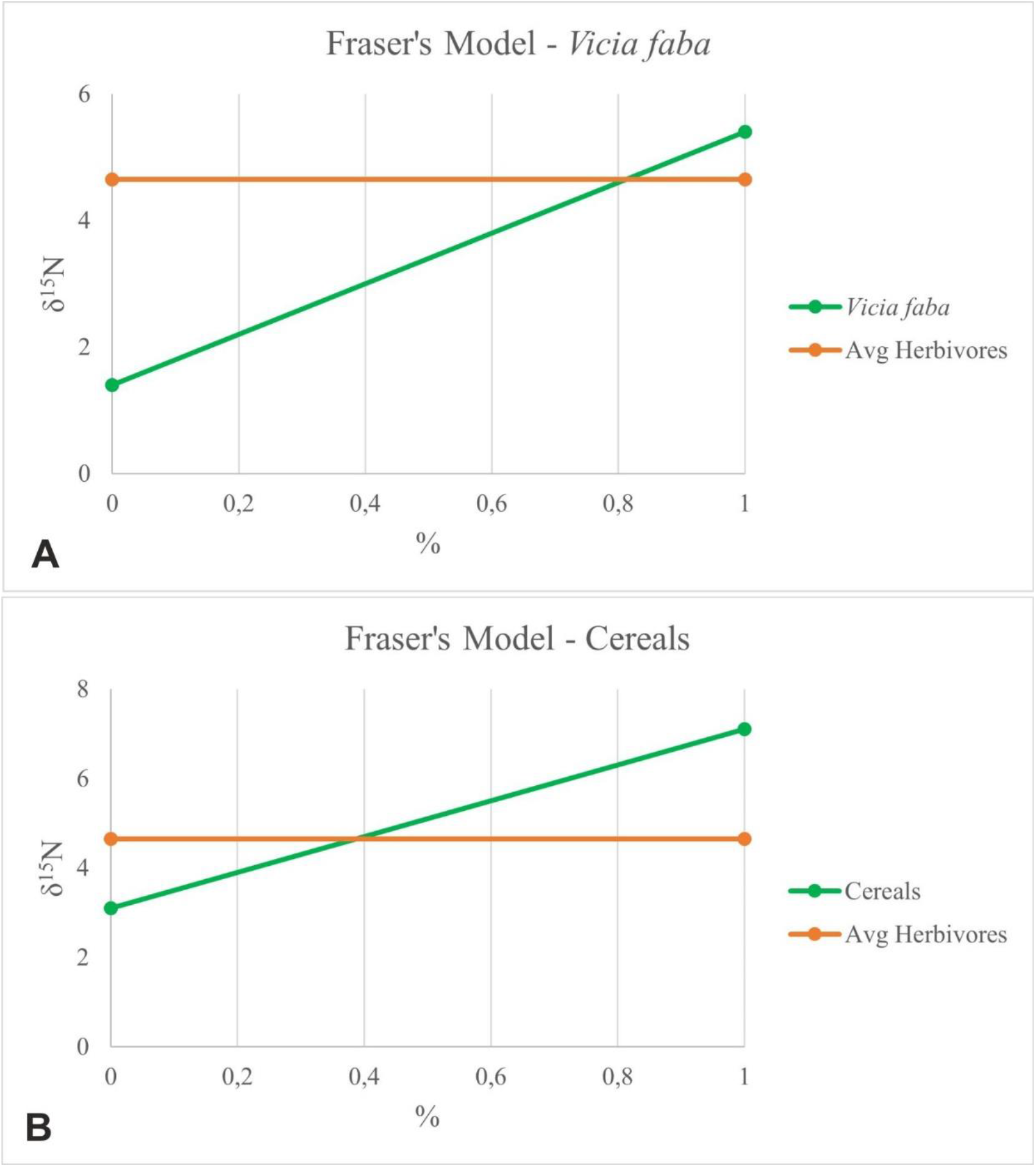
Fraser’s model (Fraser et al. 2013a). Modeled scenario estimating the plants protein fraction (percentage) from the total intake of faunal dietary choices. A) Cereals δ^15^N values determine the starting point for the esteem of the % of plants protein in faunal remains; B) Broad beans δ^15^N values determine the starting point for the estimation of the % of plants protein in the faunal remains.

We are aware that the reconstruction obtained with Fraser’s model should be interpreted with caution, considering that it reflects only the protein fraction.

Four human individuals – an adult, one skeletally immature and two children, as reported in Rolfo et al. (2021) - have been analyzed for diet reconstruction. Their values ranged between −19.9‰ and −20.7‰ (IQR= 0.39) for δ^13^C, and between 7.5‰ and 10.7‰ (IQR=1.65) for δ^15^N. Adult and young individuals have similar values, suggesting no differences in the diet despite the different ages, except for the infant of about 6 months, whose values reflect the well-known breastfeeding effects (Fuller et al. 2006; Tsutaya, Yoneda 2015). Human individual values are consistent with a mixed diet, based on terrestrial foods, more on legumes and cereals rather than on animal proteins. The consumption of freshwater fish is not excluded, while carbon values do not support any marine or C_4_ plants intake. Fraser’s model was also applied to human data to quantify the fraction of animal protein exploitation (Fig. 4). Plants and herbivores values from Pastena determine the starting point for the estimation of the percentage of animal protein in the human diet, which seems moderately high (about 70%). Still, it is essential to consider that the model is an approximation and could boost the estimation as it considers a diet exclusively (100%) based on animal protein consumption, disregarding the possible plant exploitation, even though the model prompts some initial considerations on food practices.

**Fig 4:**
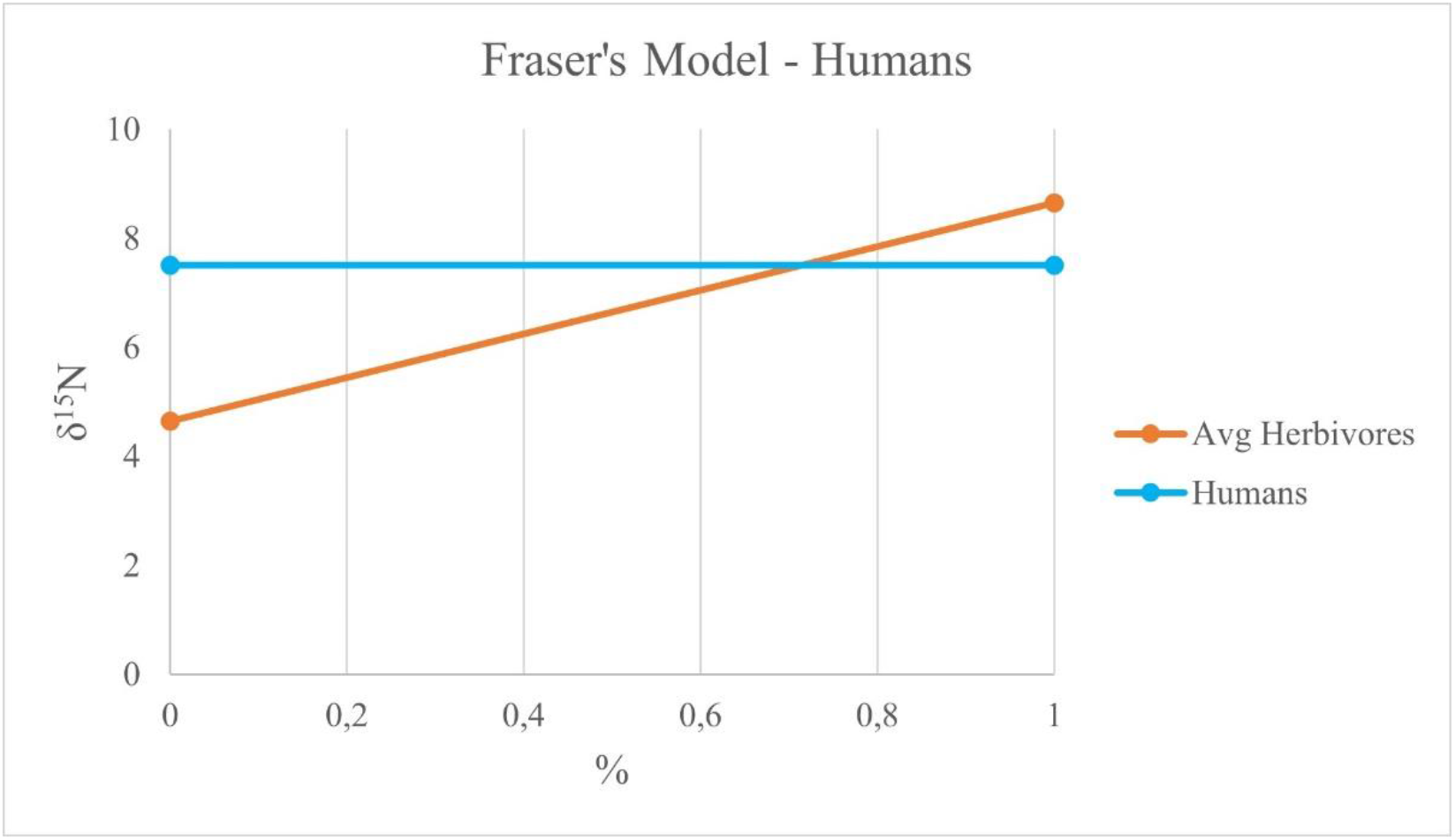
Fraser’s model (Fraser et al. 2013a). Modeled scenario estimating the protein fraction (percentage) from the total intake in the human diet. Herbivores δ^15^N determine the starting point for the estimation of the percentage of animal protein.

## Discussion

The GW2 sector of the Pastena cave is characterized by several archaeological finds, including hundreds of thousands of burnt carpological remains, mainly broad beans (*Vicia faba*). Due to the high quantity, it is possible to assume that seeds resulted from abundant production, probably due to human care and management. The amount of remains, involving remarkable energy expenditure by the human community, seems to be beyond the sole dietary consumption and could be possibly related reliably to specific rituals and, at least partially, to foraging purposes.

As previously reported (Rolfo et al. 2021), Pastena GW2 was a funerary area initially. Afterward, it appears to become devoted solely to non-burial ritual practices, as corroborated by archaeological features such as stone-paving, hearths, pits, and evidence of meals of most likely ritual nature. From that perspective, the relationship between broad beans and the after-life world seems worth mentioning. Indeed, multiple ancient cultures, such as Egyptians, Greeks, and even Romans, attested such a close link between broad beans and the soul of the dead, with the latter offering beans to keep the evil spirits away (de Cleene, Lejeune 2004; Beer 2010; Silvestri 2016).

However, the seeds should have been also part of the dietary habits of people buried inside the cave. The stable isotope data obtained from the individuals suggest a diet based principally on the consumption of C_3_ plants and the exploitation of local resources. Accordingly, we quantitatively reconstructed the human diet through a Bayesian mixed model (Fernandes et al. 2014). We implemented two different models: in the first one (Fig. 5A, Suppl. Mat.) the contribution of plants is relatively high (median 45.9% for cereals and 34% for legumes), followed by terrestrial animal meat (15.6%) and freshwater fish (4.5%). The model is consistent with the hypothesis formulated through the qualitative results obtained from stable isotope analysis and highlights the high consumption of plants and the rare assumption of freshwater fish in the community. Furthermore, considering the high number of seeds in the cave, we decided to prioritize the broad beans exploitation, even though we are aware that they could be selectively put in the cave by human selection strategy.

**Fig. 5:**
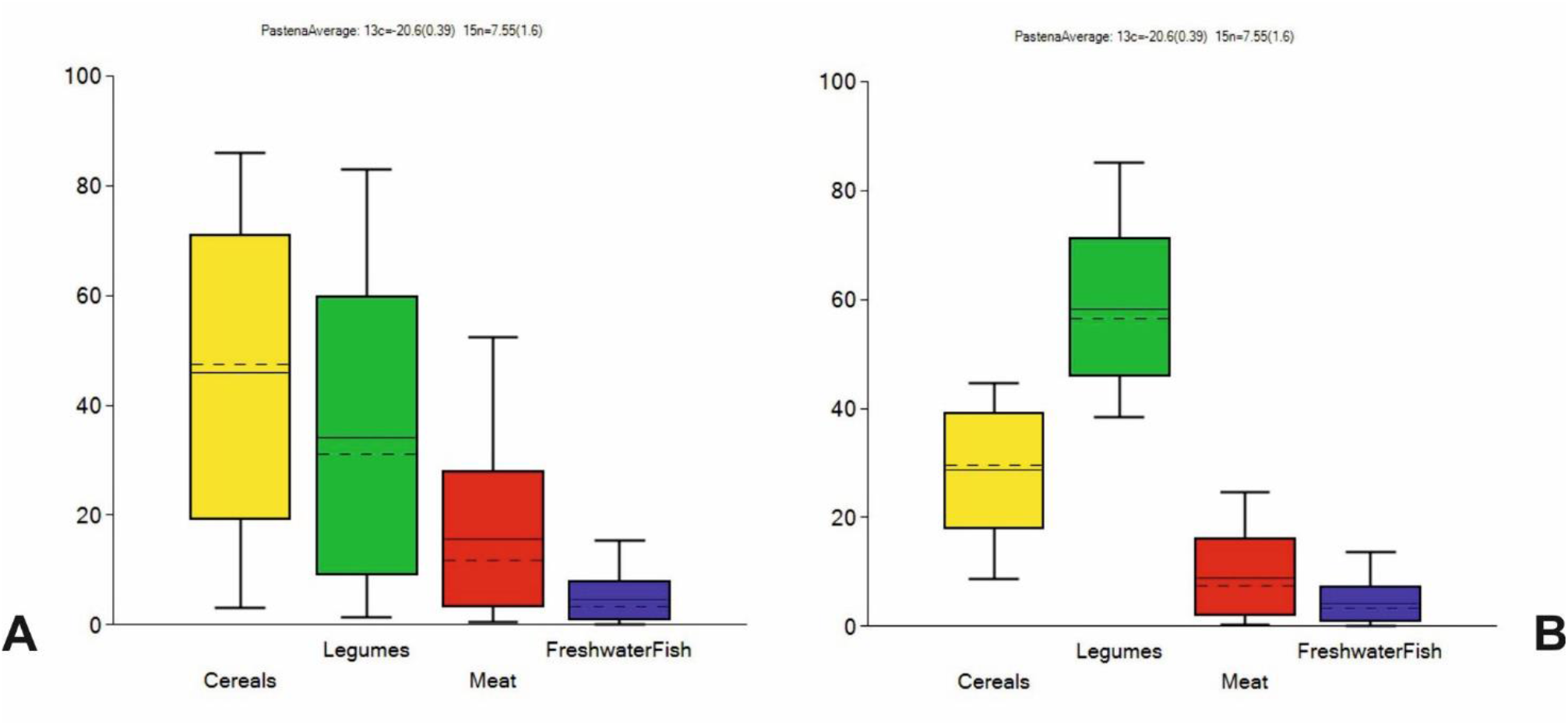
Estimated contributions of cereals, legumes, meat, freshwater fish, and secondary products to the diets of adult individuals from the Pastena cave using the Bayesian model FRUITS; A) estimations without priors; B) estimations with prioritizations (Legumes > Cereals, Cereals and Legumes > Fauna, Freshwater fish).

The prioritization of legumes and cereals led to no-significant changes in the identification of the primary edible source, even though the cereal fraction rises at the expense of the meat – (Fig. 5B, Suppl. Mat.): the intake of legumes is 58.3%, followed by cereals (28.7%), terrestrial animal meat (8.8%) and freshwater fish (4.2%).

The dietary landscape outlined for Pastena is worth being reported considering the diet reconstructions for coeval Italian sites. Only a few Central Italian sites reveal the consumption of C_4_ plants in the Middle Bronze Age (Varalli et al. 2015, Romboni et al. 2021). Genomics (Saupe et al. 2021) showed how changes occurred in central Italy during the MBA, probably related to the arrival of populations from northern areas throughout the Italian peninsula. The burial area we are considering was radiometrically dated between the end of the Early Bronze Age and the beginning of the Middle Bronze Age (Rolfo et al. 2021). Considering the geographical proximity between the Pastena cave, the Misa cave, and the La Sassa cave – where C_4_ plants consumption was identified - as well as the tight chronological frames, it is worth noting that Pastena does not seem to be involved in those complex dynamics.

Multiple factors could be responsible for that odd landscape for Pastena.

First, the slightly different chronologies of all these contexts should be considered. Indeed, the Misa cave is dated – chrono-typologically – to the MBA 1-2 (Cocchi Genick 1995), while La Sassa dates back from the Copper Age up to the MBA 3 (Alessandri et al. 2020, 2021). Remarkably, the individual from La Sassa supporting the consumption of C_4_ plants (Romboni et al. 2022) dates back to the MBA 3, which outdates the upper chronological boundary for Pastena. Additionally, the Misa cave is located in northern Latium; conversely, Pastena and La Sassa caves are both located in southern Latium and close to one another. It is worth considering that the putative demic flow responsible for introducing new crops arriving in northern Latium in the first phases of MBA might spread to southern Latium in MBA 3, excluding people buried in the Pastena cave from these complex dynamics.

Similarly to Pastena, people buried in Sepolcreto di Felcetone (northern Latium), Collepardo Cave (southern Latium), and Scoglietto cave (southern Tuscany), dated to MBA3 (Cocchi Genick 1995; Skeates et al. 2021) and EBA (Cocchi, Ceccanti 1978) respectively, did not show C_4_ consumption (Varalli et al. 2015; Skeates et al. 2021), suggesting a not continuous southward cline for the spread of the newly introduced plants as edible resources.

If people buried in the Pastena cave had a diet based on C_3_ plants, it seems reasonable to consider that they could gather these resources through the tuning of the horticultural practices. So far, the agricultural practices have been attested in Europe since the Neolithic (e.g., Buurman 1988; Fokkens 1982), and genetic studies have demonstrated how human genetic adaptation to an agricultural diet occurred during the Bronze Age (Mathieson, Mathieson 2018).

Archaeological and bioarcheological data suggest an increase in cereal cultivation in Italy, especially from the Middle Bronze Age, when intensive agriculture spread (Salvadei, Santandrea 2003; Tafuri et al. 2009; Arena et al. 2020).

Our data suggest that artificial management was applied to the growing plants. Specifically, the cereals values indicate that they grow under similar conditions. Cereals appear to be moderate-to-poorly watered (Wallace et al. 2015; Knipper et al. 2020; Varalli et al. 2021; Fig. 6A). Conversely, legumes (Fig. 6B) were well-watered (Wallace et al. 2015). However, it should be borne in mind that water availability differences might also be related to seasonal rainfall and cultivation.

**Fig. 6:**
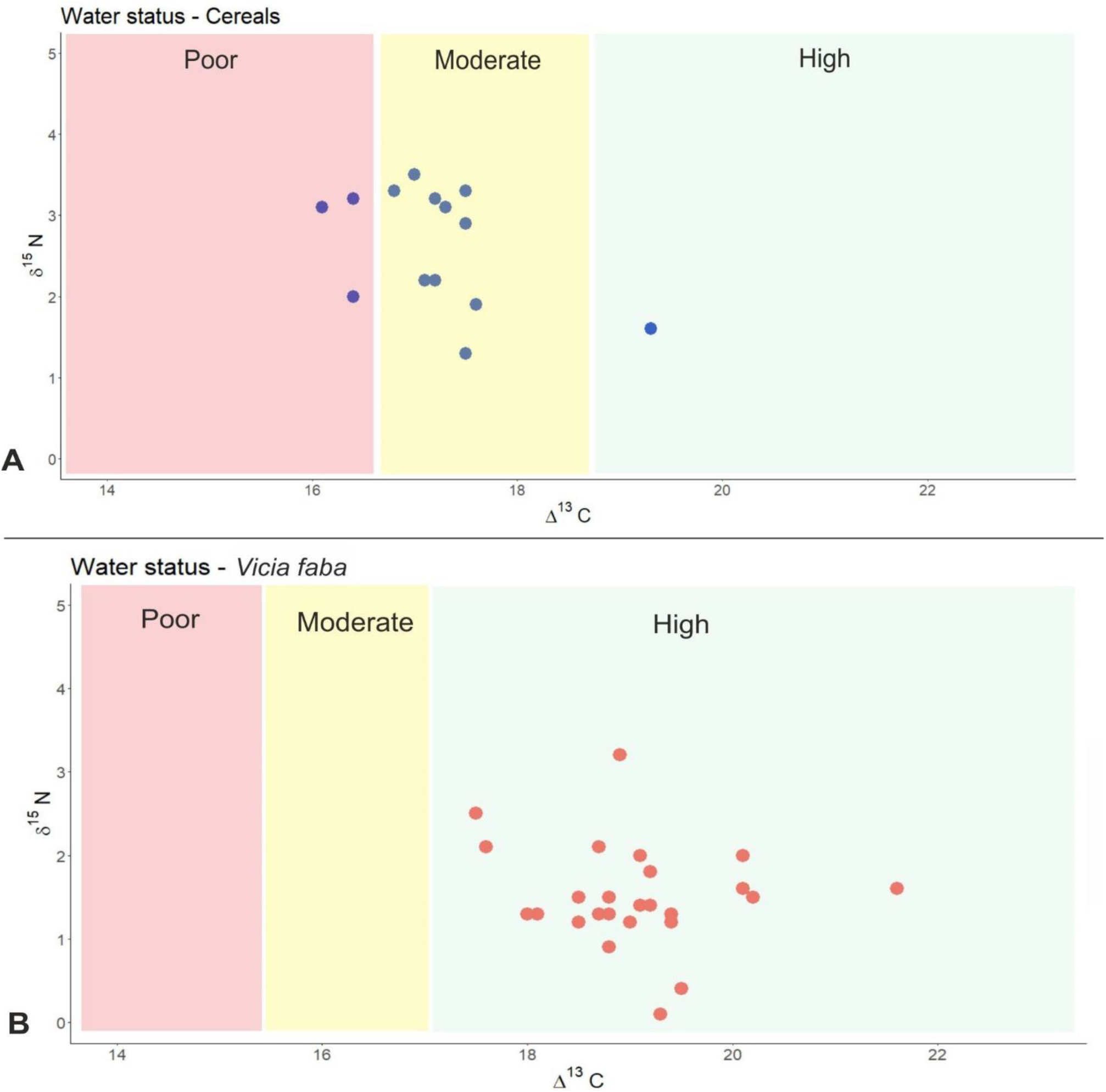
Estimations of watering status based on Wallace et al. (2015): A) Cereals; B) Broad beans

Published works (Bogaard et al. 2007, 2013; Fraser et al. 2011; Kanstrup et al. 2014) demonstrated that fertilization increases nitrogen values in cereals and legumes depending on the intensity and duration of manuring. Cereals from the Pastena cave do not appear to be extensively manured (Bogaard et al. 2013), contrarily to the legumes. Fraser et al. (2011) show that legumes have a low nitrogen isotopic signature (about 0‰), and only intensive fertilization can alter these values. *Vicia faba* δ^15^N values from the Pastena cave range from 0.1‰ to 3.2‰; thus, it is possible to hypothesize that some broad beans have been subject to fertilization. The massive presence of legumes on the site and the fact that none of them suffered from water stress led to considering these crops’ volunteer management.

Finally, it is worth mentioning that the seeds analyzed were charred, which could impact the managing identification’s reliability. However, previous studies (DeNiro, Harstoff 1985; Bogaard et al. 2007; Kanstrup et al. 2012) demonstrated that charring until 220°C does not significantly alter the pristine isotope values, which otherwise would be disrupted. We have not yet identified the temperature to which the grains were exposed; however, the influence of charring is highly questionable as some scholars claimed that the charring is not a significant modifier for δ^13^C and δ^15^N (Kanstrup 2012; Bogaard et al. 2013; Fraser et al. 2013b; Styring et al. 2013). Specifically, the carbon values were demonstrated only minimally or not affected (Fraser et al. 2013b), while δ^15^N may rise by about 1‰ (Fraser et al. 2013b). Additionally, recent research showed that the offset might be tiny and accountable by subtracting 0.31‰ from the determined values (Nitsch et al. 2015), not impacting our considerations.

## Conclusion

The GW2 sector of Pastena cave (Latium) allowed for the overall reconstruction of human subsistence strategies for one Central Italian Early-Middle Bronze Age community. Carbon and nitrogen stable isotope data confirm that people who frequented the cave were devoted to an agricultural economy, consistently with what was hypothesized through the archaeological evaluation.

The analysis performed on human remains outlines a dietary pattern based on terrestrial resources, including mainly plants and, secondarily, animal proteins.

Despite the limited sample size for humans, this paper presents, to the best of our knowledge, the first stable isotope analysis carried out on carpological remains dated to the Italian Bronze Age, contributing to filling a gap in the biomolecular research of the Italian peninsular prehistoric communities. These data allow for hypothesizing a human role in crops management. Isotope data suggest that broad beans were grown under better watering conditions than cereals, and similarly, differential fertilization can be hypothesized for these plants.

Most of the faunal remains found in GW2 belong to herbivorous and omnivorous animals, which can be left to graze, even though the relationship between crops cultivation and animal husbandry reveals that animals were probably fed, at least occasionally, with agricultural output.

Despite the limited bio-archaeological materials available in the cave, the synergic evaluation of bio-archaeological data leads to reconstructing the trophic net – from plants to humans – aiming to obtain a reliable reconstruction of the subsistence strategy for one of the communities living in an area characterized by complex demographic dynamics in Central Italian Bronze Age.

## Acknowledgments

We would thank Arturo Gnesi, Mayor of Pastena, and the community for their support since 2012, and Robin Skeates for his cooperation. The leading authors are very grateful to Marco Romboni for his support in collagen extraction.

The permits to execute the study were provided by Micaela Angle (Ministero della Cultura, Istituto Autonomo Villa Adriana e Villa d’Este) and Carlo Molle (Soprintendenza Archeologica Belle Arti e Paesaggio Frosinone e Latina). This work was supported by the Italian Ministry of Education, Universities and Research (MIUR) through PRIN 2017 action (1000 Ancient Italian Genomes: Evidence from ancient biomolecules for unravelling past human population Dynamics, GrantID: 20177PJ9XF) allotted to OR; and by a 2022 Research Fund Grant from The Prehistoric Society, Institute of Archaeology, UCL, allotted to FC. This research is part of the FC’s PhD thesis in Cultural Heritage, Education and Territory. Authors declare that they have no significant competing financial, professional, or personal interests that might have influenced the performance or presentation of the work described in this manuscript.

## Notes

### Competing Interest Statement

The authors have declared no competing interest.

